# Effects of the GluN2B antagonist, Ro 25-6981, on extinction consolidation following adolescent- or adult-onset methamphetamine self-administration in male and female rats

**DOI:** 10.1101/2020.05.11.088732

**Authors:** Sara R. Westbrook, Joshua M. Gulley

## Abstract

Previous work suggests adolescent rats have deficient extinction consolidation relative to adults. Although the mechanisms underlying this age difference are currently unknown, studies in adult rats have implicated GluN2B-containing NMDA receptor function in extinction consolidation of drug-associated memory. Importantly, GluN2B neurotransmission emerges during adolescent development, and drugs of abuse during adolescence may delay the development of extinction consolidation by disrupting the ontogeny of GluN2B function. Here, we trained Sprague-Dawley rats of both sexes to self-administer methamphetamine (METH, 0.1 mg/kg/infusion i.v.) beginning during adolescence [postnatal (P) day 41] or adulthood (P91). Rats were given short access (2 h) to self-administer METH in seven daily sessions followed by fourteen sessions with long access (6 h). Subsequently, rats underwent four daily 30-min extinction sessions with immediate post-session injections of either a GluN2B antagonist (Ro25-6981; 6 mg/kg, i.p.) or a vehicle solution. After four daily 2-h extinction sessions, a priming injection (1 mg/kg METH, i.p.) was given prior to a final 2-h reinstatement session. During LgA, adolescent-onset rats earn more METH than adult-onset rats and display greater drug-loading behavior. Rats reduced their drug-seeking behavior across extinction sessions, with no significant group differences. Rats reinstated drug-seeking following the METH priming injection, with females displaying greater reinstatement than males. These results do not support our *a priori* hypothesis that adolescent-onset METH use disrupts the ontogeny of GluN2B transmission and contributes to age-of-onset differences in extinction of METH-seeking. However, our findings suggest that age-of-onset contributes to excessive METH-taking, while sex confers vulnerability to relapse to METH-seeking.

## Introduction

Adolescent-onset drug users who develop a substance use disorder have especially high relapse rates compared to those who initiate during adulthood (Poudel and Gautam, 2017). Female users also have a higher relapse risk compared to males (Hillhouse et al., 2007; Brecht et al., 2013). The cause for this increased vulnerability to relapse in adolescents and females has not yet been uncovered, but significant effort has been placed on studying rodent models that use extinction and reinstatement paradigms. Notably, both age and sex differences have been reported in extinction, such that adolescents were impaired compared to adults in paradigms that use food or drug as reinforcers and in studies of fear conditioning (Anker and Carroll, 2010; McCallum et al., 2010; Andrzejewski et al., 2011; Zbukvic et al., 2016). Females have been shown to respond more during extinction of cocaine self-administration (Lynch & Carroll, 2000; Perry et al., 2008) and require more sessions to reduce responding during extinction of methamphetamine (METH) self-administration compared to males (Cox et al., 2013). In addition, separate studies have demonstrated that adolescent-onset and female rats exhibit greater drug-primed reinstatement than adult-onset and male rats (Anker and Carroll, 2010; Reichel et al., 2012). The mechanisms underlying this heightened relapse vulnerability in adolescents and females are not known, but a potential candidate is the delayed functional development of GluN2B-containing NMDA receptors. GluN2B neurotransmission emerges in the prefrontal cortex (PFC) during late adolescence (around P45; Flores-Barrera et al., 2014), and this developmentally-regulated process may be disrupted by drugs of abuse.

Studies using adult male rodents have implicated GluN2B receptor function in extinction of drug-seeking behavior. Non-specific blockade of NMDARs with the non-selective antagonist CPP has been shown to reduce the retention of extinction learning in rats trained to self-administer cocaine (Hafenbreidel, Todd, Twining, Tuscher, & Mueller, 2014). More selective blockade of GluN2B-containing NMDARs with ifenprodil infused directly into the infralimbic PFC has been shown to disrupt consolidation following extinction of a cocaine conditioned place preference (Otis et al., 2014). In sum, blockade of GluN2B transmission increases drug-seeking during extinction in adult males, but its role in heightened relapse risk in adolescents and females remains unclear.

Many dynamic, developmental changes are occurring in the brain during adolescence, such as in dopaminergic and glutamatergic systems in the PFC. Dopamine D_1_ receptor expression in the PFC increased to a peak at P40 before decreasing into adulthood (Andersen et al., 2000). This pruning was shown to occur specifically on PFC pyramidal cells that project to the nucleus accumbens around P44 (Brenhouse et al., 2008). The timing of this D_1_R pruning is similar to the emergence of GluN2B neurotransmission in the PFC (∼P45), which was shown to require D_1_R signaling (Flores-Barrera et al., 2014). Exposure to drugs of abuse during adolescence may detrimentally impact these developmentally regulated changes in the PFC leading to long-lasting dysfunction. Consistent with this notion, our lab has previously reported reduced D_1_R function in the mPFC (Kang et al., 2016a) and region-specific decreases in D_1_R expression in the mPFC following adolescent AMPH exposure (Kang et al., 2016b). It is not known if these drug-induced changes in D_1_Rs also impact the adolescent emergence of GluN2B signaling. However, it is notable that phosphorylation of Ser1303, which is the amino acid residue of GluN2B that is normally phosphorylated in adults following D_1_R stimulation (Sarantis et al., 2009), is decreased 24 h after 14 days of cocaine exposure in adolescent rats (Caffino et al., 2018). Taken together, these studies suggest that the reduction in D_1_Rs from adolescent drug exposure may impact the functional development of GluN2B in the mPFC, ultimately leading to impaired extinction consolidation and increased drug-seeking. However, no studies to date have tested this hypothesis as nearly all previous studies of GluN2B function in relation to heightened drug-seeking have been conducted on adults. Furthermore, the research to date has been almost exclusively limited to male subjects, preventing the direct examination of potential interactions between age and sex in conferring vulnerability to drug-seeking behavior.

The present study sought to address this gap by investigating the interaction of age and sex in drug-seeking behavior and manipulating GluN2B function during extinction consolidation to investigate the hypothesis that disrupted GluN2B functional development underlies age-of-onset differences in drug-seeking behavior. Male and female rats were trained to self-administer METH beginning in adolescence or adulthood followed by extinction sessions where drug was no longer available. GluN2B receptors were blocked with the receptor-selective antagonist Ro25-6981 during extinction consolidation to assess the role of GluN2B transmission in METH-seeking during extinction and subsequent METH-primed reinstatement. In line with previous studies of consolidation of fear conditioning (Hikind and Maroun, 2008; Sotres-Bayon et al., 2009; Otis et al., 2014), we hypothesized that GluN2B antagonism in adult-onset rats would impair extinction consolidation and increase drug-seeking behaviors, rendering their drug-seeking behavior more adolescent-like. We hypothesized that this manipulation would have minimal effects in adolescent-onset rats, but that adolescent-onset rats would display worse extinction consolidation and greater METH-seeking compared to adult-onset controls. Consistent with reports of sex differences (Lynch and Carroll, 2000; Perry et al., 2008; Cox et al., 2013), we predicted that females would exhibit greater drug-seeking than males with adolescent-onset females exhibiting the worst extinction consolidation and greatest METH-seeking behavior. All our *a priori* hypotheses, methods, and data analysis plan were pre-registered in March 2018 on the Open Science Framework (https://osf.io/avkhf).

## Methods

### Subjects

Sprague-Dawley rats (n = 60/sex), which were the offspring of 14 litters born in our facility from breeders purchased from Envigo (Indianapolis, IN), were assigned to this study. The final subject numbers were 52 male and 56 females (n = 108 rats total) as some rats were removed from the study due to loss of catheter patency (adolescent: males = 2, females = 1; adult: males = 5, females = 3) or abnormal development (1 adolescent male). Rats were housed in a temperature-controlled room on a reverse 12:12 light/dark cycle (lights off at 0900) with food and water available *ad libitum*. All experimental procedures occurred during the dark phase of the cycle. The day of birth was assigned postnatal day (P) 1 and rats were weaned into cages of two to three same-sex rats on P22. Assignment to age-of-onset groups (adolescent- or adult-onset) was balanced as best as possible within a litter. Rats were weighed daily beginning on P25. Daily checks for physical markers to estimate pubertal onset—vaginal opening in females (Castellano et al., 2011) and preputial separation in males (Korenbrot et al., 1977)—began on P30 and continued until these signs of puberty were observed in all rats. This experiment was conducted over the span of 18 months with 5 cohorts each consisting of 12 rats per age-of-onset group. All experimental procedures were in accordance with the National Research Council’s Guide for the Care and Use of Laboratory Animals and were approved by the Institutional Animal Care and Use Committee at the University of Illinois, Urbana-Champaign.

### Apparatus

Experimental sessions occurred in standard operant chambers (Coulbourn Instruments, Whitehall, PA) connected to a computer running Graphic State software (v4.1.13, Coulbourn Instruments) for automation and data collection. The operant chambers were located within ventilated, sound-attenuating cubicles. The chambers were equipped with two recessed nosepoke ports on either side of a food trough, a white cue light located above each port, a Sonalert speaker, and houselight near the ceiling on the opposite wall. An infusion pump was located outside the outer cubicle with the connected infusion line (Tygon tubing) attached to a swivel hanging from a balance arm (Instech) and protected by a metal tether inside the chamber.

### Drugs

(+) Methamphetamine HCL (METH; Sigma-Aldrich, St. Louis, MO) was dissolved in 0.9% saline and self-administered i.v. at a unit dose of 0.1 mg/kg/infusion (calculated as the weight of the salt). Thirty min prior to the reinstatement session, rats were injected with METH (1 mg/kg i.p.). Ro 25-6981, an antagonist selective for the GluN2B subunit of the NMDAR, was obtained from Hello Bio, Inc (Princeton, NJ) and dissolved in 1 part dimethyl sulfoxide (DMSO) and 2 parts 0.9% saline. Ro 25-6981 or vehicle solution was injected i.p. immediately after each of the four short extinction sessions at a dose of 6 mg/kg (1 ml/kg volume).

### Surgery

Starting the day prior to catheterization surgery, rats were given the antibiotic Trimethoprim Sulfa (1.1% v/v; Midwest Veterinary Supply) in their drinking water with solutions changed every 2-3 days. On the day of surgery (P32 ± 2 days for adolescent-onset or P82 ± 2 days for adult-onset), rats were implanted with an indwelling catheter in the right jugular vein as described previously (Hankosky et al., 2018). In brief, the right jugular vein was isolated under isoflurane anesthesia (2-4% Midwest Veterinary Supply) and a small incision was made on the vein to insert the catheter. Once inserted, the remaining portion of the catheter was tunneled subcutaneously over the shoulder where it was externalized via an access port in the mid-scapular area. Rats were allowed at least 5 days of recovery from surgery before beginning self-administration sessions. Catheters were flushed daily with 0.1 ml of 50 U/mL heparinized saline to assist with maintaining catheter patency. Patency was assessed once before self-administration procedures began, after the final self-administration session, and as often as once a week if patency loss was suspected. These checks were conducted by flushing the catheter with 0.07-0.1 ml 15% ketamine (100 mg/mL) and 15% midazolam (5 mg/mL) solution with rapid loss of muscle tone a positive indicator of catheter patency.

### Self-Administration

The self-administration procedures were the same as previously described (Westbrook et al., 2020). On P40 ± 2 days for adolescent-onset or P90 ± 2 days for adult-onset, rats were placed in the operant chambers for a 90-min habituation session during which the nosepoke ports were covered. The next day rats began 2-h short access (ShA) self-administration sessions wherein a response in the active port (always the right port) resulted in an infusion of 40-80 μL METH (2-4 s duration, dependent on body weight) delivered in a unit dose of 0.1 mg/kg/inf. Illumination of the white cue light above the active port and a 2.9 kHz, 80 dB tone occurred for 4 s concurrent with METH infusion onset. A 20-s timeout period, wherein responses into the active port had no programmed consequences, was signaled by illumination of red lights inside both nosepoke ports. Rats received 7 daily 2-h ShA sessions followed by 14 daily 6-h long access (LgA) sessions. LgA sessions were the same as ShA sessions except for their increased length and the addition of a cap on the maximum number of infusions set to 120 (12 mg/kg/session) in order to prevent accidental overdose.

### Extinction and Reinstatement

One day following the final LgA session, rats underwent four daily 30-min extinction sessions wherein nosepoke responses in the previously reinforced port had no programmed consequence. Immediately following these sessions, rats received a challenge injection of the GluN2B antagonist, Ro25-6981 (6 mg/kg, i.p.), or an equivalent volume of vehicle (1ml/kg, i.p.). This dose was chosen based on previous studies showing impairment of behavioral flexibility and fear extinction in adult rats (Dalton et al., 2008, 2011, 2012). For the next 4 days, rats received daily 2-h sessions to assess extinction retention. Similar procedures have been used previously with cocaine self-administration to demonstrate the effects of infralimbic prefrontal cortex inactivation on extinction consolidation (LaLumiere et al., 2010) and in studies of impairment and facilitation of extinction consolidation following systemic administration of an NMDAR antagonist and a partial agonist, respectively (Hafenbreidel et al., 2014). The following day, a priming injection of METH (1 mg/kg, i.p.) was administered 30-min prior to a 2-h session under extinction conditions. This dose was based on a previous study examining sex differences in adult rats (Reichel et al., 2012). Responding during this METH-primed session was compared to responding during the last extinction session as a measure of reinstatement of METH-seeking.

### Data Analysis

#### Pre-registered Analyses

These analyses were planned *a priori* and detailed in the pre-registration of this study (https://osf.io/avkhf). Nosepokes into the reinforced and unreinforced ports, as well as infusions earned were measured. The number of infusions earned was multiplied by the unit dose (0.1 mg/kg/infusion) to calculate METH intake (mg/kg). Daily METH intake and first hour METH intake were used as the dependent variables in self-administration analyses. Between-subjects factors included sex, age-of-onset, and treatment; the within-subjects factor was session. Separate three-way mixed ANOVAs (sex x age-of-onset x session) were used to assess METH intake during ShA and LgA sessions. Escalation was assessed via separate one-way repeated measures ANOVAs within each sex and age-of-onset group with Tukey post-hoc comparisons of METH intake in each LgA session compared to the first LgA session. Cumulative METH intake (mg/kg) was also calculated for all self-administration sessions and this measure was used as a covariate in the analyses of extinction and reinstatement responding.

Nosepokes into the previously reinforced port during short extinction sessions (sessions 1-4) were analyzed using mixed ANCOVAs with sex, age-of-onset, treatment, and session as factors. To assess extinction consolidation, mixed ANCOVA (sex x age-of-onset x treatment x session) was conducted on previously reinforced nosepokes during the long extinction sessions (sessions 5-8). Differences in previously reinforced nosepokes during the first 30 min of extinction session 5 were determined via three-way ANCOVA with sex, age-of-onset, and treatment as factors. Previously reinforced nosepokes during the last extinction session and the METH-primed reinstatement session were analyzed using a mixed ANCOVA with the factors of sex, age-of-onset, treatment, and session. Finally, comparison of previously reinforced nosepokes between the final extinction session and reinstatement session was conducted within each group using pre-planned Tukey post-hoc tests.

#### Exploratory Analyses

Additional analyses, which were not pre-planned, were performed to compare the results of this study to those in our recent work (Westbrook et al., 2020) demonstrating age-of-onset differences in self-administration behavior. First, body weight was analyzed across the peri-adolescent period (P25-P74) using separate two-way mixed ANOVAs (age-of-onset x postnatal day) for each sex. Second, further analysis of escalation was conducted on METH intake during the first and last LgA sessions using a three-way mixed ANOVA (sex x age-of-onset x session). Lastly, during our pre-planned analysis of extinction responding, it was noted that there was no significant effect of session after accounting for the relationship of the covariate (cumulative METH intake). Therefore, exploratory analysis of previously reinforced nosepokes was assessed in the first 30 min of these sessions—the same duration as the first four extinction sessions—to determine if rats extinguished their previously reinforced response across sessions. This analysis was performed using a four-way mixed ANCOVA (sex x age-of-onset x treatment x session) with cumulative METH intake as the covariate.

## Results

### METH Self-administration

#### Body Weight

Analysis of body weight from P25 to P74 demonstrated the expected weight gain across postnatal days in *ad libitum* fed Sprague-Dawley rats (Fig. 2). However, adolescent-onset males had suppressed weight gain relative to their adult-onset counterparts that remained in their homecages during this time [Age-of-onset by postnatal day interaction: F(49,2450)=3.15, *p*<0.0001]. Post-hoc analyses indicated that this METH-induced suppression of weight gain began on P47 near the start of LgA sessions and persisted through P74 (all *p*’s<0.01). In females, adolescent-onset and adult-onset groups did not significantly differ on any postnatal day in post-hoc tests, despite a significant age-of-onset by postnatal day interaction [F(49,2646)=1.85, *p*=0.0003]. Thus, the weight gain suppressing effect of METH was only present in adolescent-onset males.

**Figure 1.**
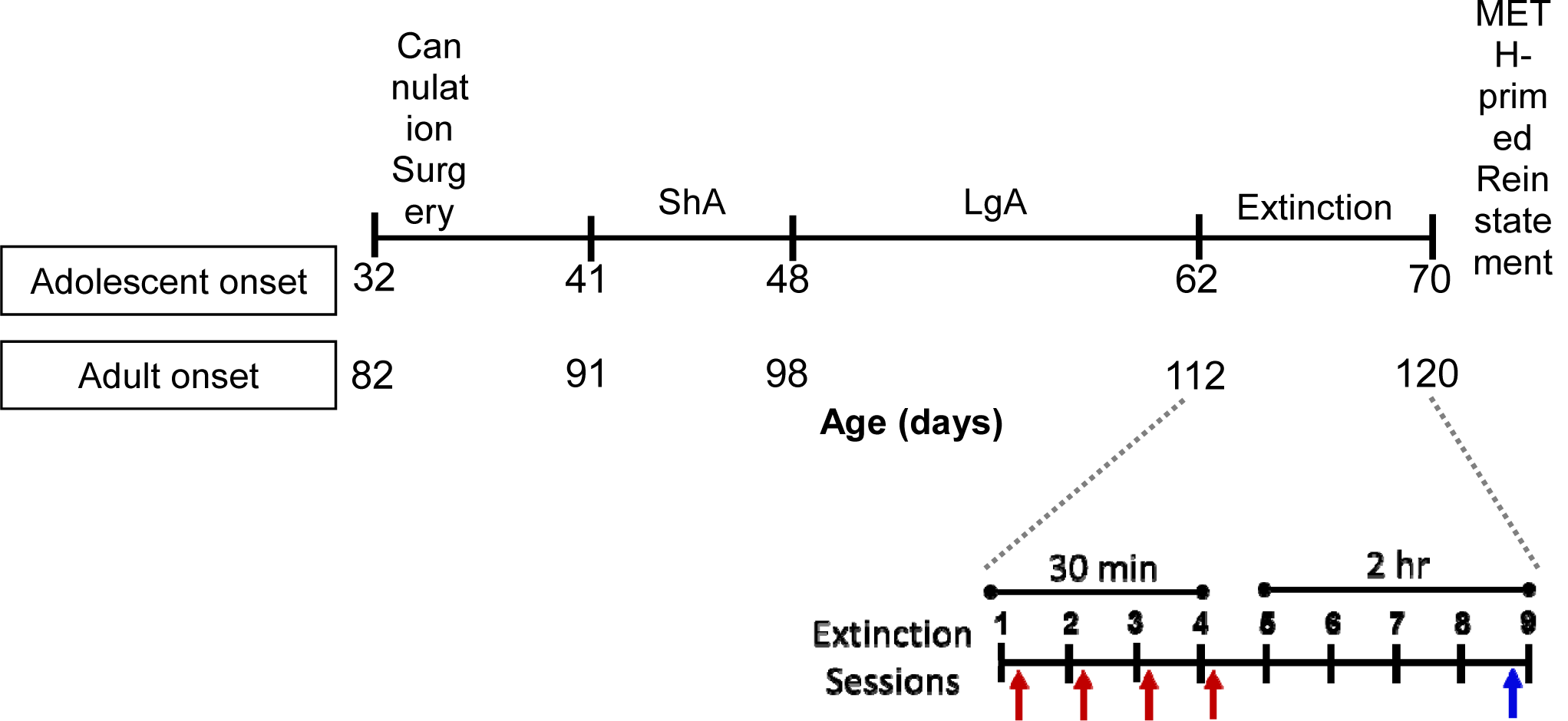
Experimental timeline for rats that underwent short access (ShA) followed by long access (LgA) METH self-administration. The injection schedule for the extinction sessions and METH-primed reinstatement is shown in more detail. Arrows indicate the timing of i.p. injections: red = 6 mg/kg Ro25-6981 or vehicle given after the session; blue = 1 mg/kg METH given before the session

**Figure 2.**
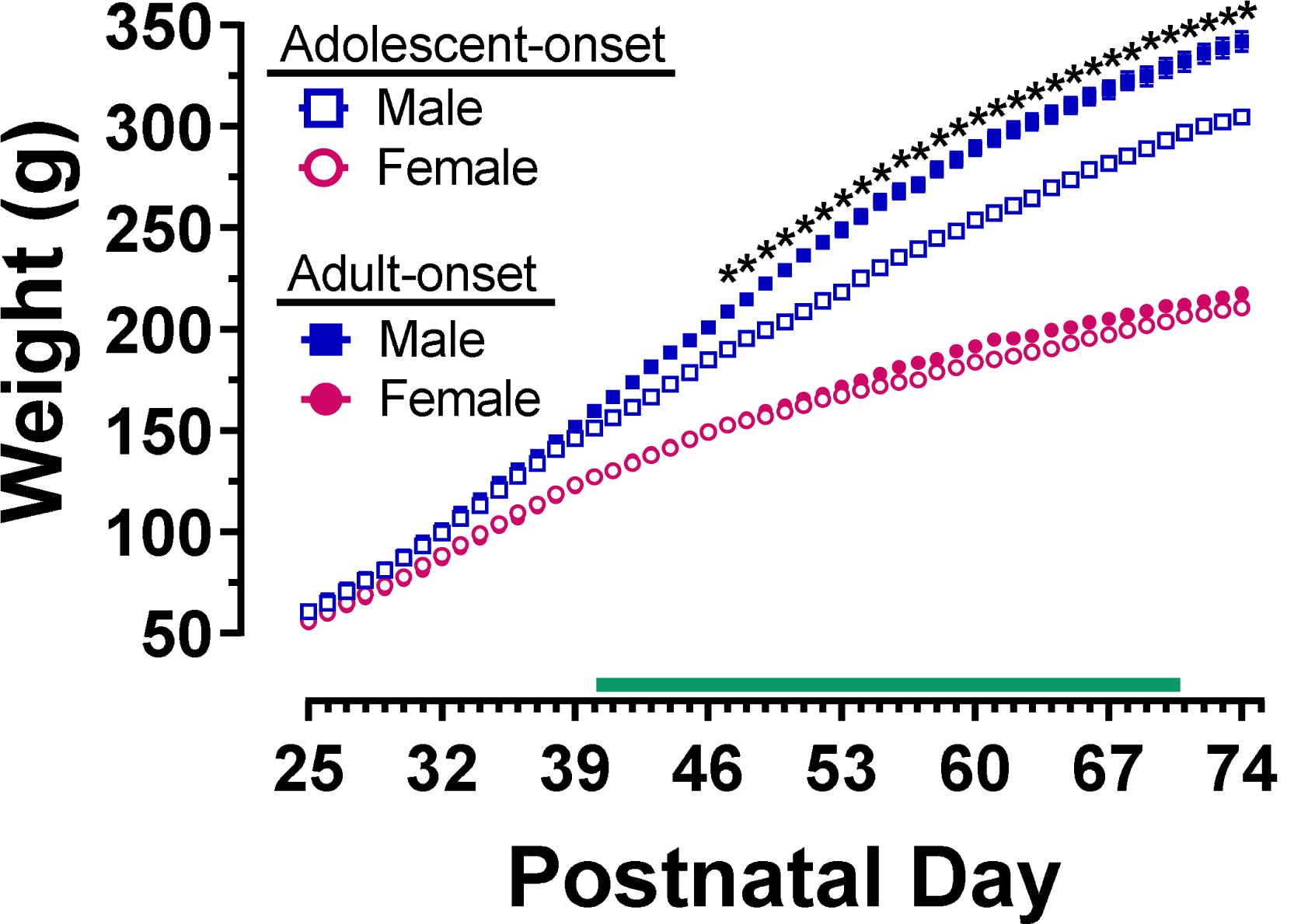
Body weights (g) for rats that began METH self-administration sessions during adolescence or adulthood. The horizontal green bar indicates the time period when adolescent-onset rats were performing experimental sessions. During this time, their adult-onset counterparts remained in their homecages. \**p* < 0.05 vs. adolescent-onset males on the noted days

#### Short Access

Three-way mixed ANOVA conducted on METH intake during ShA sessions (Fig. 3A) indicated a significant main effect of session [F(6,104)=25.03, *p*<0.0001] and an interaction of sex and session [F(6,104)=2.56, *p*=0.0237]. METH intake on the first session was significantly greater than subsequent ShA sessions (all *p*’s<0.0001). However, post-hoc tests did not reveal significant sex differences for any of the ShA sessions.

**Figure 3.**
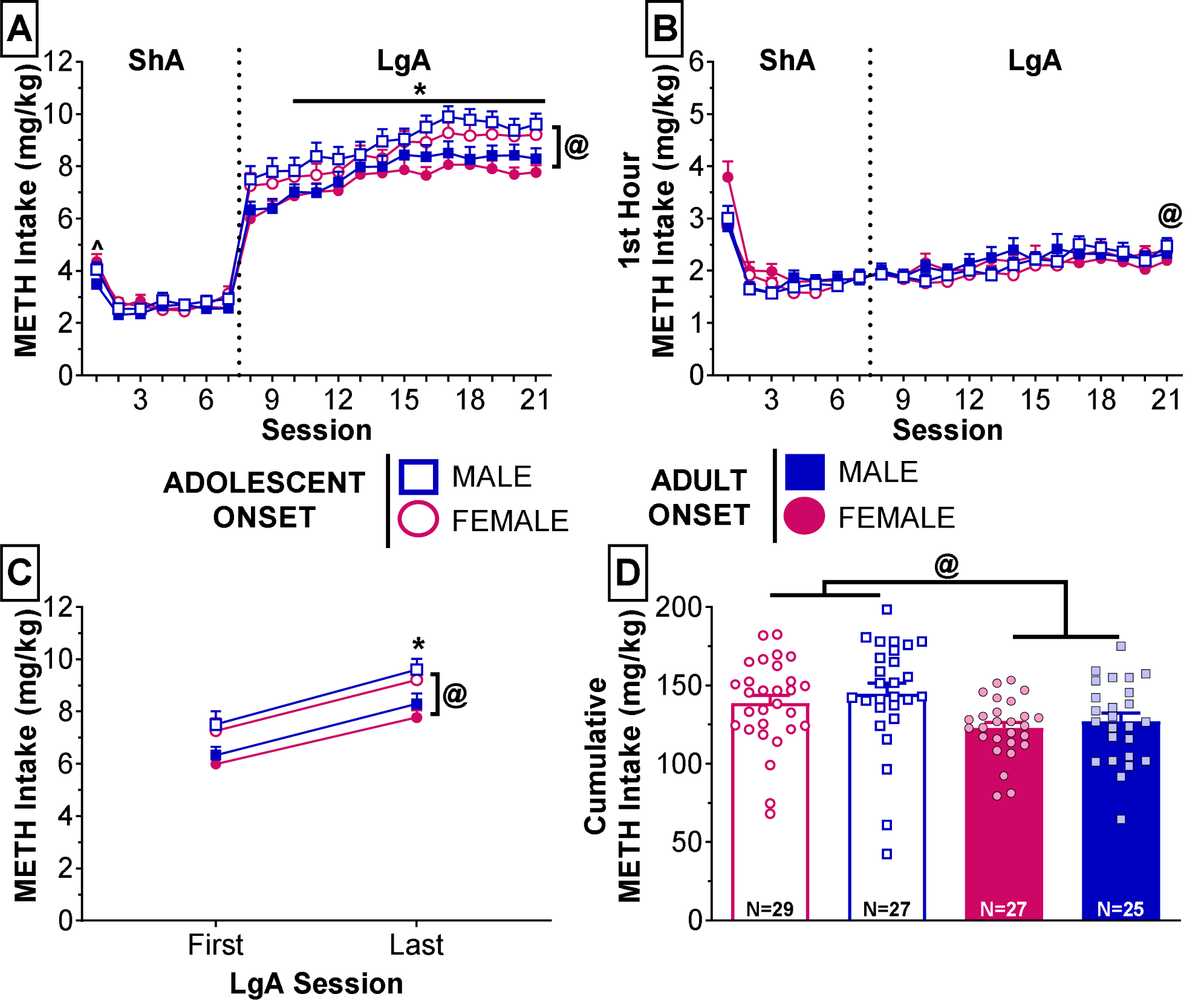
METH self-administration that began during adolescence or adulthood in rats of both sexes. (**A**) METH intake (mg/kg) during 7 ShA (2-h) sessions followed by 14 LgA (6-h) sessions. \**p* < 0.05 vs. session 8 (first LgA session) collapsed across sex and age-of-onset, ^@^*p* < 0.01 collapsed across sex and session. (**B**) METH intake (mg/kg) during the first hour of the daily self-administration sessions. ^@^*p* < 0.01 vs. session 8 (first LgA session) for adolescent-onset rats collapsed across sex. (**C**) METH intake (mg/kg) during the first and last LgA sessions. \**p* < 0.05 vs. session 8 (first LgA session) collapsed across sex and age-of-onset, ^@^*p* < 0.01 collapsed across sex and session. (**D**) Cumulative METH intake (mg/kg) summed across all self-administration sessions. ^@^*p* < 0.01 collapsed across sex

#### Long Access

During LgA sessions (Fig. 3A), adolescent-onset rats earned more METH than adult-onset rats [main effect of age-of-onset: F(1,104)=10.07, *p*=0.0020]. METH intake was significantly escalated relative to the first LgA session’s intake [main effect of session: F(13,104)=10.14, *p*<0.0001]. This effect was apparent starting on session 10 (*p*=0.0114) and remained escalated through the rest of the LgA sessions (all *p*’s<0.01). One-way repeated measures ANOVAs revealed significant effects of session for each group [adolescent-onset: females, F(13,364)=2.44, *p*=0.0035, males, F(13,338)=2.89, *p*=0.0006; adult-onset: females, F(13,336)=2.47, *p*=0.0032, males, F(13,312)=2.72, *p*=0.0012]. Adolescent-onset females and males began escalating their METH intake on session 15 (*p*=0.0214) and 14 (*p*=0.0479), respectively, while adult-onset females and males started escalating on session 13 (females: *p*=0.0030, males: *p*=0.0066).

METH intake during the first hour of LgA sessions was also assessed with a three-way mixed ANOVA, which revealed a significant main effect of session [F(13,104)=5.05, *p*<0.0001] and a significant age-of-onset by session interaction [F(13,104)=2.63, *p*=0.0032]. Adolescent-onset rats, but not adult-onset rats, demonstrated significant escalation of their first hour METH intake, specifically on the last LgA session (*p*=0.0411); adolescent- and adult-onset groups did not significantly differ from each other on any LgA session. One-way repeated measures ANOVAs within each group showed significant main effects of session for each group, except adolescent-onset females [adolescent-onset males: F(13,26)=7.95, *p*<0.0001; adult-onset: females F(13,26)=4.89, *p*=0.0003, males F(13,312)=2.09, *p*=0.0148]. Notably, post-hoc tests comparing each LgA session to the first hour intake of the first LgA session only revealed significant escalation of first hour intake in adolescent-onset males on the last LgA session (*p*=0.0451).

Exploratory analysis of escalation during LgA sessions revealed that rats significantly escalated their METH intake from the first to the last LgA session (Fig. 3C; main effect of session [F(1,104)=85.49, *p*<0.0001]). Adolescent-onset rats had significantly higher METH intake than adult-onset rats [main effect of age-of-onset: F(1,104)=15.61, *p*=0.0001]. Finally, age-of-onset differences in measures of escalation of METH intake across sessions was further reflected in cumulative METH intake (Fig. 3D) across all self-administration sessions. Adolescent-onset rats earned higher cumulative METH intake than adult-onset rats [main effect of age-of-onset: F(1,104)=9.78, *p*=0.0023]. There were no significant sex differences or interaction of age-of-onset and sex.

### Extinction and Reinstatement

#### Extinction

Nosepoke responses into the previously reinforced port were analyzed using 4-way ANCOVA with cumulative METH intake as the covariate (Fig. 4). During the first four extinction sessions, the relationship of the covariate and nosepokes depended on group and session [Table 1; 5-way interaction with covariate: F(3,92)=3.04, *p*=0.0327]. After accounting for cumulative METH intake, there were no remaining significant main effects or interactions. For the last four extinction sessions, the relationship between cumulative METH intake and previously reinforced nosepokes varied by session [Table 2; F(3,92)=3.50, *p*=0.0186]. After accounting for cumulative METH intake, there were no significant main effects or interactions. We also assessed extinction consolidation by analyzing previously reinforced responses during the first 30 min of extinction session 5. Four-way ANCOVA revealed no significant main effects or interactions after accounting for the covariate. However, the relationship of cumulative METH intake and previously reinforced responding did vary significantly depending on sex and drug challenge [Table 3; F(1,92)=5.43, *p*=0.0220]. The positive correlation only reached statistical significance for vehicle-treated females.

**Table 1.** Linear regressions of 5-way interaction with cumulative intake for previously reinforced nosepokes measured during the first four extinction sessions.

| Session | Age | Sex | Trt | Intercept | Slope | r<sup>2</sup> | p |
| --- | --- | --- | --- | --- | --- | --- | --- |
| 1 | Adolescent-onset | F | Ro | -24.741 | 0.41632 | 0.20799 | 0.08753 |
|  |  |  | Veh | 4.683 | 0.1156 | 0.09139 | 0.29349 |
|  |  | M | Ro | 0.07 | 0.1355 | 0.13529 | 0.21629 |
|  |  |  | Veh | 14.347 | 0.05939 | 0.14126 | 0.18538 |
|  | Adult-onset | F | Ro | 31.329 | -0.11212 | 0.01663 | 0.66044 |
|  |  |  | Veh | 2.718 | 0.1321 | 0.2562 | 0.07757 |
|  |  | M | Ro | -16.568 | 0.29032 | 0.42716 | **0.02117\*** |
|  |  |  | Veh | -1.15 | 0.15057 | 0.09267 | 0.31189 |
| 2 | Adolescent-onset | F | Ro | -20.032 | 0.28189 | 0.39214 | **0.01251\*** |
|  |  |  | Veh | -2.599 | 0.1223 | 0.20621 | 0.10287 |
|  |  | M | Ro | 10.719 | 0.00547 | 0.00063 | 0.93488 |
|  |  |  | Veh | 11.589 | 0.00755 | 0.0036 | 0.83862 |
|  | Adult-onset | F | Ro | 1.178 | 0.13363 | 0.03952 | 0.49569 |
|  |  |  | Veh | 1.694 | 0.10982 | 0.11378 | 0.2597 |
|  |  | M | Ro | 8.693 | 0.04919 | 0.03075 | 0.58569 |
|  |  |  | Veh | -10.828 | 0.22876 | 0.35676 | **0.03112\*** |
| 3 | Adolescent-onset | F | Ro | -12.646 | 0.19912 | 0.19589 | 0.09852 |
|  |  |  | Veh | -6.584 | 0.14133 | 0.30299 | **0.04138\*** |
|  |  | M | Ro | 1.744 | 0.06533 | 0.08307 | 0.33958 |
|  |  |  | Veh | 3.03 | 0.06422 | 0.07036 | 0.35939 |
|  | Adult-onset | F | Ro | 18.071 | -0.02042 | 0.00166 | 0.88989 |
|  |  |  | Veh | 0.97 | 0.08953 | 0.15024 | 0.19066 |
|  |  | M | Ro | -8.92 | 0.19101 | 0.37594 | **0.034\*** |
|  |  |  | Veh | -10.118 | 0.17803 | 0.51291 | **0.00589\*** |
| 4 | Adolescent-onset | F | Ro | -4.95 | 0.15867 | 0.07186 | 0.33406 |
|  |  |  | Veh | -4.223 | 0.08646 | 0.36844 | **0.02134\*** |
|  |  | M | Ro | 5.336 | 0.04548 | 0.02951 | 0.57469 |
|  |  |  | Veh | 10.728 | 0.01524 | 0.00955 | 0.73968 |
|  | Adult-onset | F | Ro | -16.569 | 0.22921 | 0.33783 | **0.02926\*** |
|  |  |  | Veh | -6.608 | 0.13161 | 0.4146 | **0.01755\*** |
|  |  | M | Ro | 5.574 | 0.03893 | 0.04302 | 0.51774 |
|  |  |  | Veh | -23.678 | 0.29385 | 0.58484 | **0.00233\*** |

\***p**<0.05
Ro = Ro 25-6981
Veh = vehicle

**Table 2.** Linear regressions of 2-way interaction with cumulative intake for previously reinforced nosepokes during the last four extinction sessions.

| Session | Intercept | Slope | $r^2$ | $p$ |
| --- | --- | --- | --- | --- |
| 5 | 7.3149 | 0.1482 | 0.06962 | **0.00579\*** |
| 6 | 2.5143 | 0.1248 | 0.06833 | **0.00628\*** |
| 7 | 8.7421 | 0.0471 | 0.01805 | 0.16565 |
| 8 | 1.7735 | 0.0853 | 0.0583 | **0.01182\*** |

**\* $p < 0.05$**

**Table 3.** Linear regressions of 3-way interaction with cumulative intake for previously reinforced nosepokes during the first 30 min of extinction session 5.

| Sex | Trt | Intercept | Slope | $r^2$ | $p$ |
| --- | --- | --- | --- | --- | --- |
| F | Ro | 22.1985 | -0.07266 | 0.04907 | 0.24816 |
| F | Veh | -16.6906 | 0.20707 | 0.34448 | **0.00129\*** |
| M | Ro | 6.1769 | 0.02583 | 0.01549 | 0.55336 |
| M | Veh | 5.7748 | 0.02796 | 0.02139 | 0.46666 |

**\* $p < 0.05$**
Ro = Ro 25-6981
Veh = vehicle

**Figure 4.**
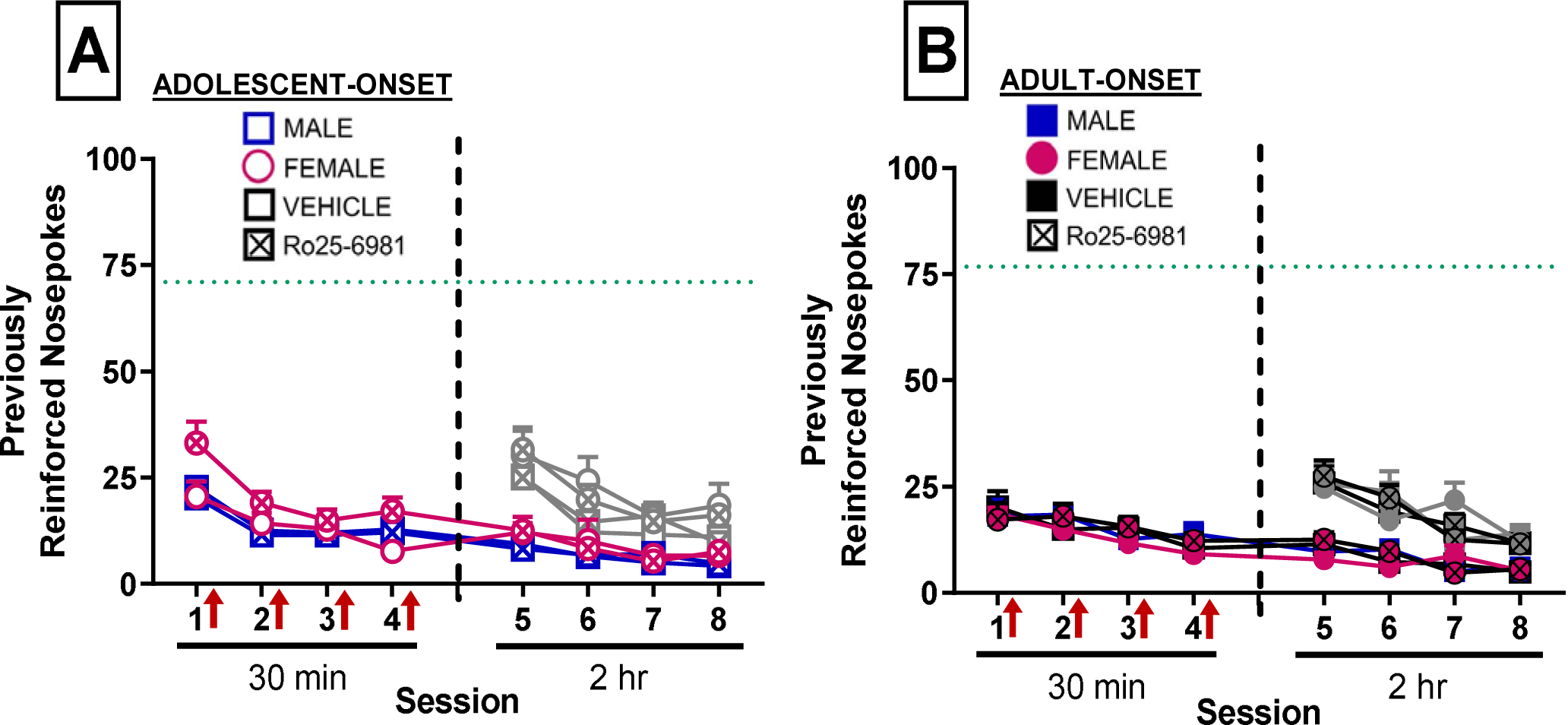
Responses into the previously reinforced nosepoke port during sessions under extinction conditions. For 2-h sessions, responding during the first 30 min is shown in line with the previous four sessions that were each 30 min in duration; gray symbols represent responding during the entire 2-h session. Green dotted lines indicate average nosepokes collapsed across groups during the first 2 hours of the final LgA session. (Adolescent-onset: n = 13-15/group; Adult-onset: n = 12-14/group)

Exploratory analysis of the first 30 min of extinction sessions 5-8 revealed that the positive correlation between cumulative METH intake and previously reinforced nosepokes depended on drug challenge and session [Table 4; F(3,92)=2.91, *p*=0.0385]. After accounting for the covariate, a significant drug challenge by session interaction remained [F(3,92)=3.14, *p*=0.0292]. Post-hoc tests did not reveal significant differences between drug challenge groups for any of the sessions, although both drug challenge groups significantly reduced their previously reinforced responses across sessions.

**Table 4.** Linear regressions of 3-way interaction with cumulative intake for previously reinforced nosepokes during the first 30 min of extinction sessions 5-8.

| Session | Trt | Intercept | Slope | $r^2$ | $p$ |
| --- | --- | --- | --- | --- | --- |
| 5 | Ro | 12.5986 | -0.00988 | 0.00151 | 0.78012 |
|  | Veh | -4.6679 | 0.10993 | 0.15677 | **0.00304\*** |
| 6 | Ro | 1.8716 | 0.04605 | 0.0597 | 0.07497 |
|  | Veh | -1.1272 | 0.07162 | 0.04266 | 0.13402 |
| 7 | Ro | -1.0974 | 0.05197 | 0.07152 | 0.05059 |
|  | Veh | -0.2207 | 0.04995 | 0.06019 | 0.07377 |
| 8 | Ro | 0.006 | 0.04273 | 0.09877 | **0.02064\*** |
|  | Veh | 1.4507 | 0.03134 | 0.0654 | 0.06198 |

\* **$p < 0.05$**
Ro = Ro 25-6981
Veh = vehicle

#### Reinstatement

Previously reinforced responses following a METH-priming injection were compared to responses during the final extinction session with a four-way ANCOVA (covariate: cumulative METH intake). After accounting for the significant positive relationship of cumulative METH intake and previously reinforced nosepokes [r^2^=0.0261, *p*=0.0174; F(1,100)=7.51, *p*=0.0073], a significant sex by session interaction remained [F(1,99)=4.62, *p*=0.0340]. Males and females increased their previously reinforced responding (Fig. 5) during the METH-primed session compared to the final extinction session (*p*’s<0.0001), but females reinstated their responding to a greater extent than males (*p*=0.0024). Reinstatement was also assessed within each group via pre-planned comparisons between sessions. Adolescent-onset females challenged with Ro25-6981 or the vehicle solution significantly reinstated their previously reinforced responding (*p<*0.0001 and *p*=0.0116, respectively). Adolescent-onset males challenged with Ro25-6981 significantly increased their responses (*p*=0.0282). Reinstatement of responding was also significant in adult-onset females challenged with the vehicle solution (*p*=0.0082). The remaining groups—adolescent-onset males challenged with vehicle, adult-onset males regardless of drug challenge, and adult-onset females challenged with Ro25-6981—failed to reach statistical significance in the pre-planned comparisons.

**Figure 5.**
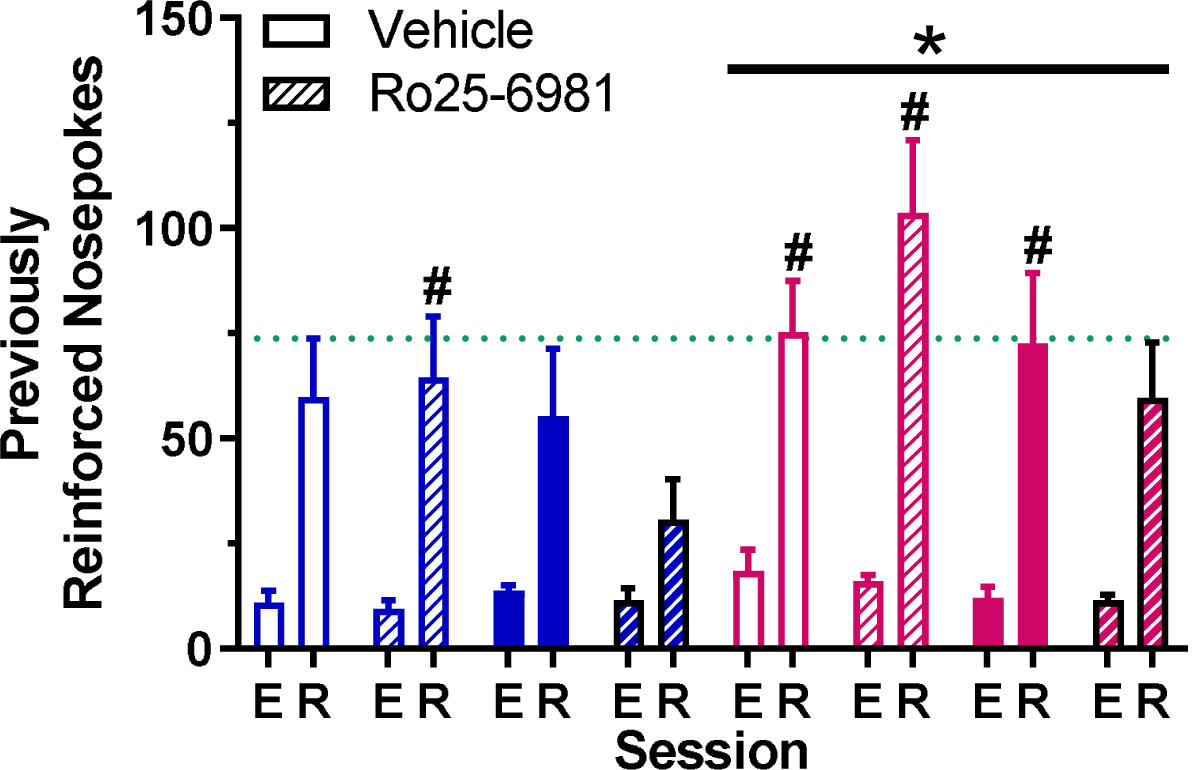
Previously reinforced nosepoke responses during the final extinction session (E) compared to the METH-primed reinstatement (R) session. Green dotted line indicates average nosepokes collapsed across groups during the first 2 hours of the final LgA session. \**p* < 0.01 vs. males for the reinstatement session collapsed across age-of-onset and drug challenge group, ^#^*p* < 0.05 vs. last extinction session within group. (Adolescent-onset: n = 13-15/group; Adult-onset: n = 12-14/group)

## Discussion

GluN2B-containing NMDA receptor transmission emerges during adolescence, which may contribute to the development of adult-like extinction consolidation. Disruption of this developmentally regulated event by drugs of abuse may lead to enhanced drug-seeking behaviors, contributing to adolescents’ heightened relapse rates. The goal of the present study was to investigate the potential role of GluN2B function in age-of-onset and sex differences in extinction consolidation and subsequent reinstatement following METH self-administration. We found that adolescent-onset rats have higher METH intake than adult-onset rats under long access conditions. Post-session GluN2B blockade did not impact drug-seeking during extinction, and we found no age-of-onset or sex differences during extinction. In response to a METH priming injection, female rats, regardless of age-of-onset and previous drug challenge, reinstated their drug-seeking to a greater extent compared to male rats. Our hypothesis that disrupted GluN2B function contributes to age-of-onset and sex differences in drug-seeking during extinction was not supported here. However, our findings do suggest that adolescent-onset rats may more readily lose control of their drug-taking, while females may be more susceptible to relapse above and beyond what would be predicted from their previous cumulative drug intake.

During ShA, METH intake for all groups was relatively stable except for the first session. Increased intake on the first session likely occurs because the nosepoke ports were made accessible for the first time and were thus more salient compared to the previous habituation session where the ports were covered. METH intake during subsequent sessions was consistent with previous work reporting stable intake during ShA and age differences in escalation of intake during LgA (Anker et al., 2012; Westbrook et al., 2020). Adolescent-onset rats took more METH than adult-onset rats throughout the LgA sessions, including significant escalation of intake in the first hour of the final LgA session, consistent with our recent report (Westbrook et al., 2020). This escalated METH intake may indicate that adolescent-onset rats have developed habitual or compulsive drug-taking. Previous work has demonstrated that in rats given prolonged access to self-administer cocaine, drug-seeking is resistant to suppression by the presence of an aversive conditioned stimulus, suggesting that the drug-seeking behavior has become compulsive (Vanderschuren and Everitt, 2004; Pelloux et al., 2007). Moreover, extended cocaine intake, as opposed to extended conditioning factors, was shown to be sufficient for the development of this compulsive drug-seeking (Jonkman et al., 2012). Interpreting our current study’s findings as a greater susceptibility to develop compulsive METH-taking in adolescent-onset rats compared to adult-onset rats would be consistent with previous reports of adolescents exhibiting greater drug-taking in the face of negative consequences, specifically aversive histamine injections (Holtz and Carroll, 2015), as well as continued drug-seeking during periods of non-availability (Anker et al., 2011) compared to adult rats. However, further research is needed using METH self-administration with punishment conditions, such as footshock or histamine injections, in order to directly test whether the greater escalation of METH intake in adolescent-onset compared to adult-onset rats is indicative of the development of a compulsion to self-administer METH.

In the present study, we did not find evidence for sex differences in escalation during LgA. This lack of effect is not consistent with our previous report (Westbrook et al., 2020) or others using METH (Reichel et al., 2012) or cocaine (Algallal et al., 2019). In our previous study, the effect of sex that we reported appeared to be primarily driven by adolescent-onset females. The discrepancy in sex differences between our studies may be partly attributed to individual differences. In our first report, our adolescent-onset female group of 13 rats showed the greatest variability in LgA intake of all our groups. In contrast, all our groups in the current study have comparable variability that is reduced relative to our previous study due to the larger sample sizes (n=25-29). Therefore, it is possible that our previously reported sex effect driven by adolescent-onset females may have resulted from the sample containing a higher proportion of individuals with increased escalation. Neither of the studies from our lab, nor a recent report from another lab (Daiwile et al., 2019), are consistent with two studies that found adult females escalated their drug intake to a greater extent than adult males (Reichel et al., 2012; Algallal et al., 2019). This difference could be due to animal housing procedures between labs. Westenbroek and colleagues (2013) found that isolated females take more cocaine and escalate their motivation to respond for cocaine more than their isolated male counterparts, and pair-housing attenuates these effects in females. Individual housing may contribute to females’ greater escalation of METH (Reichel et al., 2012) and cocaine intake (Algallal et al., 2019), while our lab and the Cadet lab group-housed the rats and did not observe greater escalation in females (Daiwile et al., 2019; Westbrook et al., 2020).

In the current study, we did not find evidence for any group differences in responding during extinction sessions following adolescent- or adult-onset METH self-administration. Previous reports of adolescent deficits in extinction consolidation were primarily studied in fear conditioning paradigms (for review, see Baker, Den, Graham, & Richardson, 2014); however there are some reports of greater responding in adolescents compared to adults during extinction of instrumental responding for food (Andrzejewski et al., 2011) or drug reinforcers (Anker and Carroll, 2010). The absence of an age-of-onset effect in our study may be due to the ages we tested or the housing conditions. In Anker and Carroll’s study, the adolescent group began cocaine self-administration between P26 and P28 and concluded around 10 days later when extinction began. In our study, adolescent rats self-administered METH from P41-62. In the fear extinction literature, adolescent deficits in extinction consolidation have been shown to occur around P35 (McCallum et al., 2010; Kim et al., 2011). If there is a relatively short window of vulnerability for drugs of abuse to prevent the development of adult-like extinction consolidation, then it is possible that our study may have started self-administration after that window had passed while the Carroll lab’s study targeted this window. We selected the adolescent age of P41 for self-administration in our study in order to coincide with the adolescent developmental changes in D_1_R expression and GluN2B function (Andersen et al., 2000; Brenhouse et al., 2008; Flores-Barrera et al., 2014) as our hypothesis was that METH’s disruption of these specific developmental events underlies heightened drug-seeking behaviors in adolescent-onset rats. However, it may be the case that drugs of abuse may have a greater impact on extinction behavior if taken earlier in adolescence.

During the first four days of extinction, rats received systemic injections of a GluN2B-selective antagonist or vehicle solution. This drug challenge was given post-session, as opposed to before the session, in order to target the consolidation of the recent extinction learning that occurred in the short extinction sessions (LaLumiere et al., 2010; Hafenbreidel et al., 2014). The drug challenge did not appear to affect drug-seeking in any of our groups. One possibility is that the dose we selected for our antagonist challenge was not effective; however, this is unlikely given that we chose this dose based on previous literature showing its significant effects on fear extinction and behavioral flexibility in adult male rats (Dalton et al., 2008, 2011, 2012). Moreover, this dose was shown to attenuate LTP and block LTD in vivo in the hippocampus with these effects measured up to 2 hours after the i.p. injection (Fox et al., 2006). The time course for the onset of Ro 25-6981’s behavioral effects is not entirely clear, but we do know that it can impact behavior 30 min post-injection (Dalton et al., 2008, 2011, 2012) and the functional effects in the brain can last up to 2 hours (Fox et al., 2006). The timeline of Ro 25-6981’s functional effects is likely within the range for impacting consolidation of extinction memory, which was found to be no longer NMDA receptor dependent 2 hours after the extinction session (Burgos-Robles et al., 2007). However, it is important to note that it is possible that our drug treatment missed the window for effect, if consolidation of the extinction memory occurred very rapidly (<30 min post-session) and Ro 25-6981’s onset of action was no shorter than 30 mins.

Another potential explanation for the lack of effect of GluN2B antagonism on extinction consolidation is that the role of GluN2B transmission in consolidation may depend on the type of extinction—Pavlovian vs. instrumental. In adult male rodents, extinction of Pavlovian fear conditioning was disrupted by systemic GluN2B antagonism (Dalton et al., 2012) and intra-infralimbic GluN2B blockade impaired extinction consolidation of environmental cues in a cocaine conditioned place preference paradigm (Otis et al., 2014). In contrast, GluN2B blockade did not impact extinction of instrumental responding for a drug reinforcer in the present study or another study (Hafenbreidel et al., 2017).

Following extinction sessions, rats underwent METH-primed reinstatement. Rats reinstated their drug-seeking response during the reinstatement session relative to the final extinction session and this reinstated drug-seeking behavior was positively correlated with cumulative METH intake during self-administration. Females reinstated their drug-seeking response to a greater extent compared to males consistent with the previous literature (Reichel et al., 2012; Cox et al., 2013; Ruda-Kucerova et al., 2015), and this sex effect was still evident after controlling for the positive relationship of cumulative METH intake on reinstated drug-seeking. Since we did not find group differences in performance during extinction sessions, our finding of sex differences in reinstatement cannot be attributed to differential extinction learning or memory. Our assessment of METH-primed reinstatement occurred on a single day, so it is possible that the estrous cycle could impact our findings and contribute to the observed sex difference. We did not monitor estrous cycle in our females because it is unclear how to control for exposure to this invasive procedure in male rats. Moreover, it is unlikely that estrous cycle phase fully explains the sex difference in our study because previous studies have failed to demonstrate a significant effect of estrous cycle phase on measures of METH self-administration, extinction, and METH-primed reinstatement (Reichel et al., 2012; Cox et al., 2013; Ruda-Kucerova et al., 2015).

Our finding of a sex effect in reinstatement was not influenced by previous drug challenge or age-of-onset. Previous examination of age differences in cocaine-primed reinstatement have found that adolescent heightened reinstatement is dose-dependent (Anker and Carroll, 2010). We selected our dose for the METH priming injection based on the dose-response curve for sex differences in METH-primed reinstatement conducted by Reichel and colleagues (2012). We chose the 1 mg/kg dose because it produced the most pronounced sex difference in adult rats; however, this dose may not be the most appropriate dose for observing age-of-onset differences in reinstatement. Future work using additional doses is needed to determine whether age-of-onset differences in METH-primed reinstatement are dose-dependent like is seen with cocaine-primed reinstatement.

In conclusion, our findings do not support the notion that disruption of GluN2B functional development impairs extinction of the instrumental drug-seeking response to contribute to heightened drug-seeking behaviors. Adolescent-onset rats may more readily lose control of their METH intake, which may partly explain this population’s heightened vulnerability to developing problems with drug abuse. Greater METH intake during self-administration predicted greater reinstatement of drug-seeking. Although females did not have greater METH intake compared to males during self-administration, females still displayed greater reinstatement. This sex effect above and beyond the relationship of cumulative METH intake on later drug-seeking, suggests that there are other mechanisms that underlie female vulnerability to relapse besides drug intake. Future investigations are needed to examine the underlying mechanisms of adolescent enhanced susceptibility to lose control of drug-taking and increased vulnerability to relapse in females.

## Acknowledgements

The authors thank Erika Carlson, Kate Hamblen, Kailey Komnick, Karen Lai, Jacob O’Russa, Brittany Rhed, Tugba Serbest, Arch Topouzian, and Julia Wisowaty for excellent technical assistance.

